# AutoGater: A Weakly Supervised Neural Network Model to Gate Cells in Flow Cytometric Analyses

**DOI:** 10.1101/2022.12.07.519491

**Authors:** Mohammed Eslami, Robert C. Moseley, Hamed Eramian, Daniel Bryce, Steven B. Haase

## Abstract

Flow cytometry is a useful and efficient method for the rapid characterization of a cell population based on the optical and fluorescence properties of individual cells. Ideally, the cell population would consist of only healthy viable cells as dead cells can confound the analysis. Thus, separating out healthy cells from dying and dead cells, and any potential debris, is an important first step in analysis of flow cytometry data. While gating of debris can be conducted using measured optical properties, identifying dead and dying cells often requires utilizing fluorescent stains (e.g. Sytox, a nucleic acid stain that stains cells with compromised cell membranes) to identify cells that should be excluded from downstream analyses. These stains prolong the experimental preparation process and use a flow cytometer’s fluorescence channels that could otherwise be used to measure additional fluorescent markers within the cells (e.g. reporter proteins). Here we outline a stain-free method for identifying viable cells for downstream processing by gating cells that are dying or dead. AutoGater is a weakly supervised deep learning model that can separate healthy populations from unhealthy and dead populations using only light-scatter channels. In addition, AutoGater harmonizes different measurements of dead cells such as Sytox and CFUs.

## Introduction

Flow cytometers provide single-cell based measurements of components that are fluorescently tagged. They are now critical instruments for a range of cell biology applications from synthetic biology, immunology, and biomarker discovery^1^. In these applications, informative cells are generally alive and in cases where dead or dying cells compose a substantial fraction of the population, methods are needed to identify dead cells so they can be removed from the analysis by gating.

There are a variety of tools that support the gating process from manual, expert driven selection of events^2^, to data-driven based clustering methods ^3–9^. Recently, there has been a considerable amount of research in using neural network models to identify cell populations of interest ^10–14^. While these new models are trained with the abundance of data present in flow cytometry, they do not directly encode information about each event for the model. Instead, they rely on sample information, such as disease not-disease, and show that filters learned can correlate with known phenotypic signatures (such as CD8+ cells).

This training problem is further complicated when the phenotype of interest does not have a clear, consolidated definition, such as live and dead cells. Current methods of differentiation between live and dead yeast cells mainly rely on adding stains to cells prior to usage of a flow cytometer. There are two stain types that are commonly used, both of which rely on the fact that cells with membrane damage usually die^15^. The first type of stain can only penetrate into cells that have broken membranes, allowing for dead or damaged cells with ruptured membranes to be identified ^15^. These kinds of stains include Sytox ^16^, propidium iodide, ethidium bromide, trypan blue, or erythrosine B ^15^. The second type of stain can penetrate live cells as well, but are able to be pumped out or reduced by living cells while dead cells cannot do the same ^15^. Examples of these stains include phloxine B or methylene blue ^15^. Another stain can be used to measure mitochondrial function where dead cells are defined by lack of mitochondrial function rather than a lack of membrane integrity or pump function. These staining methods allow researchers to distinguish between live and dead cells at an individual cell level, but they can be time consuming, expensive, and add complexity to the experiment. In addition, usage of stains requires one of the flow cytometer’s fluorescence channels to be dedicated to measuring stain presence, thus making that channel unavailable for other experimental measurements and/or create additional needs for controls and fluorescence compensation when spectral overlaps exist.

In some cases, live and dead cells can be differentiated growing cells on solid or liquid medium ^15^. Perhaps the most common of these methods is Colony Forming Units (CFUs) ^15^. While using CFUs is considered to be a “gold standard”, there are two main limitations to this method. First, CFUs can only provide a **population-based** estimate of the percentage of live and dead cells, and cannot determine if any **individual** cell is live or dead ^15^. CFUs do not measure cell death directly, but rather provide a growth inhibition measure that can be used as a proxy to measure cell death. In doing so, they fail to capture some viable yeast cells such as those who remain alive but are unable to divide ^15^. Thus, CFUs cannot distinguish between cytostatic and cytotoxic impacts and may overestimate the percentage of dead cells if some proportion of the cells remain alive but are unable to divide (cell-cycle arrest) ^15^.

Here, we present AutoGater, a weakly-supervised neural network model trained with a novel objective function to support identification of dead cells without the use of stains (Figure 1). AutoGater’s objective harmonizes event-based and population-based notions of death into a single model to improve learning from events by corroborating events with population data. Experiments were performed on budding yeast cells (*Saccharomyces cerevisiae*) to weakly label events from a flow cytometer as alive or dead. A two stage-model was then trained to label events from all other conditions as alive or dead. The model was validated from experiments that used two different modes of killing, ethanol and heat, and across time course measurements. The model’s predictions were compared to both event based methods using stains as well as population based measurements in the form of CFUs.

**Figure 1.**
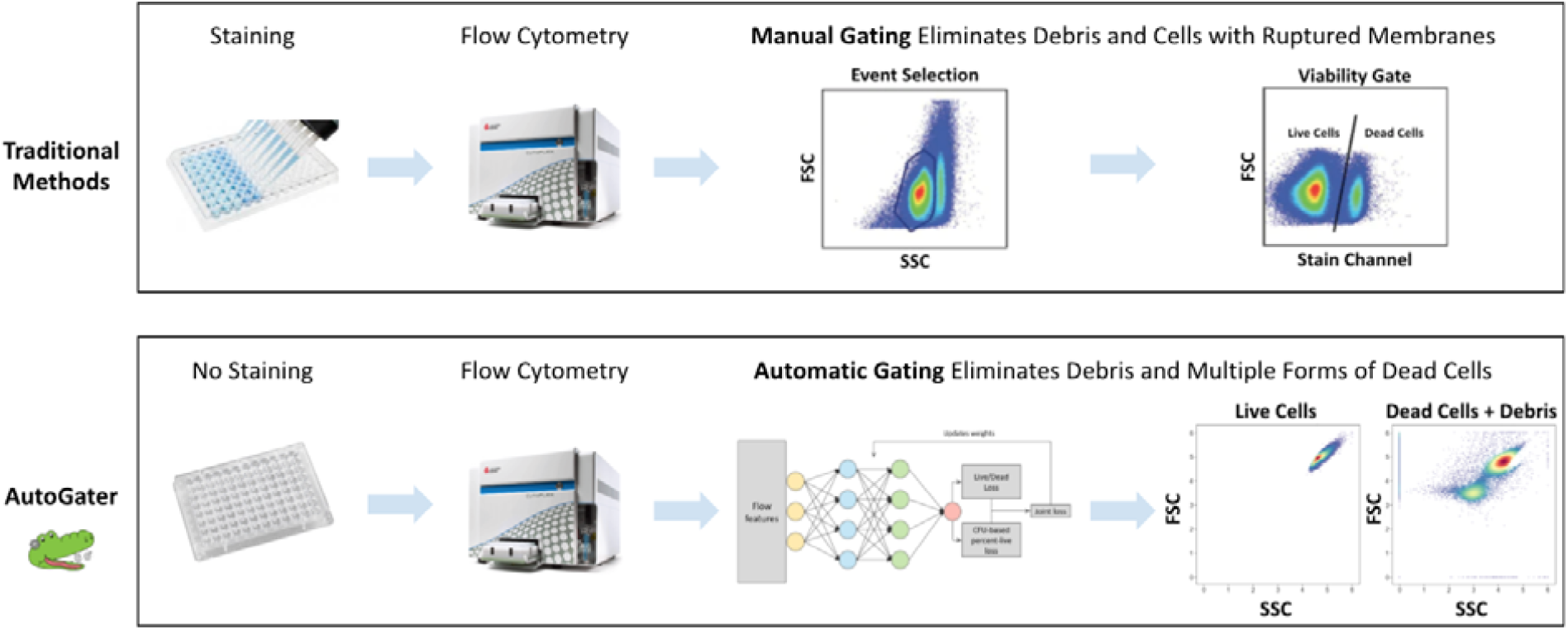
Overview of the AutoGater methodology and its benefits. Manual Gating figures are from ^17^.

## Methods

### Experiments to Generate Flow Cytometry and CFU Training Data

All experiments used the S288C wild-type strain of *S. cerevisiae*. In order to train the model, we treated cell populations to two stressful conditions that are capable of killing cells in the population: high ethanol levels and high temperature. We varied the level of these stressors in hopes that we would see increasing levels of cell death within the population as the stress was increased. Before treatment with various ethanol concentrations or temperatures, yeast cells were grown in rich YEP medium (1% yeast extract, 2% peptone, 0.012% adenine, 0.006% uracil) containing 2% dextrose (YEPD) at a temperature of 30°C until a cell density of ∼3e6 cells/mL was reached. Upon reaching the required cell density, cells were either treated with various ethanol concentrations (0%, 5%, 10%, 12%, 15%, 20%, and 80%) or temperatures (25°C, 30°C, 35°C, 40°C, 45°C, 50°C, 55°C, and 65°C). Ethanol treated cells were kept at a temperature of 30°C.

The sampling regime for ethanol treated cells was 0, 0.5, 3 and 6 hours, while the sampling regime for temperature treated cells was 0, 0.5, 1, 2, 3, 4, 5, and 6 hours. The time course and gradient of treatments resulted in 28 and 64 different conditions for ethanol and temperature, respectively. Effectively, this created a gradient consisting of 28 and 64 samples with potentially different levels of cell death in the population, which we believe should capture both dead and dying yeast cells.

In an effort to compare multiple methods for assessing cell death, each sample was split into 3 samples: one for inoculating a plate with ∼300 cells for subsequent CFU counting, one for staining with SYTOX Orange Nucleic Acid Stain (Invitrogen) and the last kept untreated with SYTOX. The last two samples, one with SYTOX Orange added and one without it added, were then run through an Attune NxT Flow Cytometer (ThermoFisher Scientific) with 488 nm and 561 nm lasers and fluorescent protein filter kit. The YL1 detector with the 585/16 band pass filter was used in the acquisition of SYTOX Orange fluorescence.

### Two Stage Training of Neural Network with Custom Objective Function

The neural network model to gate cells goes through two stages of training. The first stage learns weights from a set of weak labels on events, while the second stage updates the weights based on information from CFU assay (Figure 2B). Another description of the two stages is that the first stage provides a set of preliminary labels to events that are then modified based on information from the additional assay.

**Figure 2:**
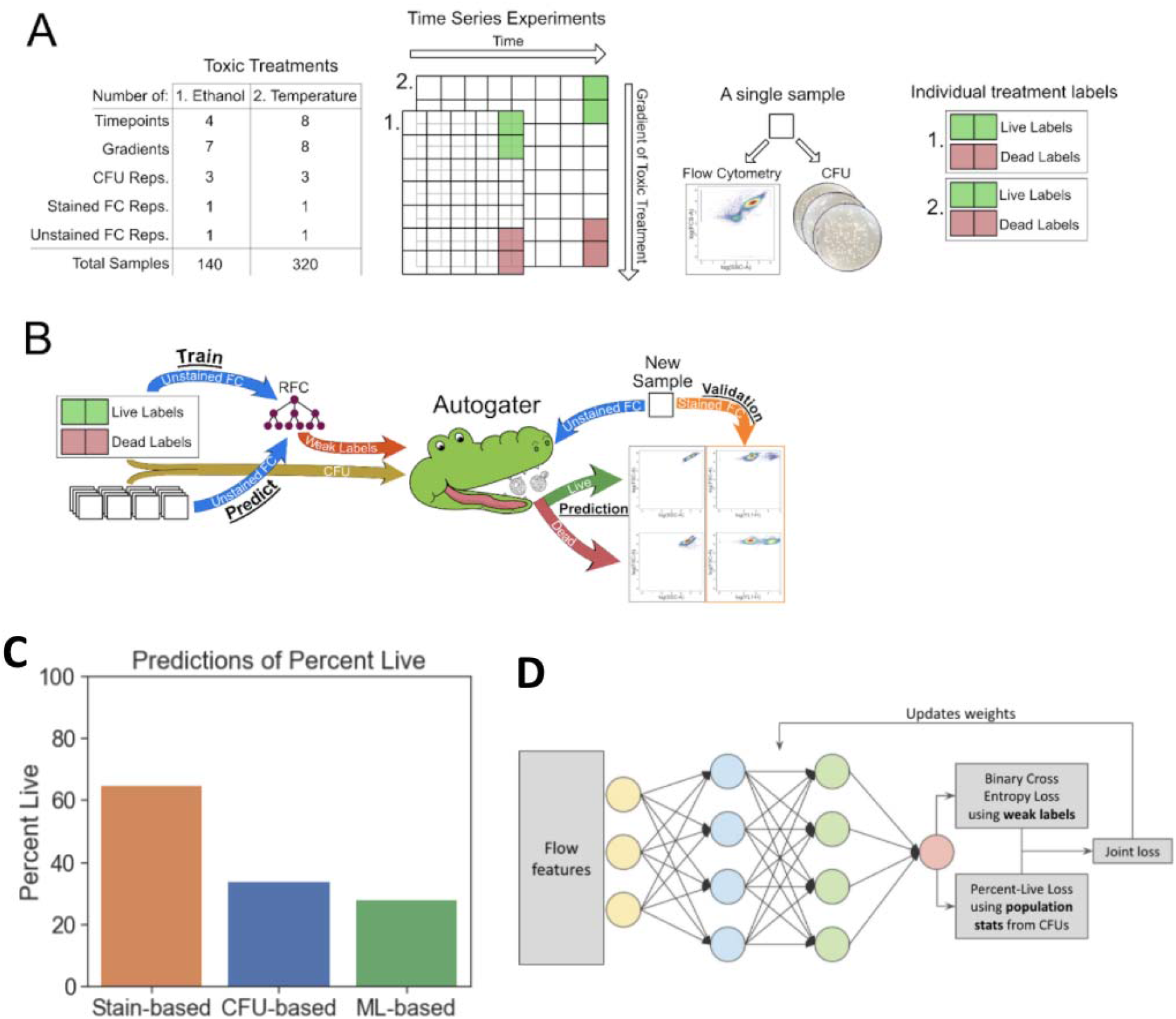
Autogater labeling and training framework. A) Gradients of two modes of killing (heat and ethanol) were used to generate data for training. Four conditions were used for training and all remaining were used for test. B) Autogater’s two stage framework first predicts the set of held out conditions based on the weak labels using a random forest classifier (RFC), and then adjusts those predictions based on the CFU data at that condition using a neural network. C) The three different methods provide three very different assessments of percentages of live cells for the same sample. Methods are needed to harmonize across these different assessments. D) Architecture of a neural network that takes non-color channels from a flow cytometer as input (FSC, SSC) and is trained to jointly optimize cell-based and population based notions of death.

#### Stage 1: Weakly Supervised Model for Annotating Dead Cells

The first stage of training relies on a set of weak labels attached to every event. In the case of the live/dead phenotype, we select a subset of experimental conditions to serve as “live” and “dead” conditions. Most of the 28 and 64 different conditions for ethanol and temperature contained a mixture of live and dead cells making this labeling effort by hand impossible. However, a subset of the conditions did contain mostly, if not all, live or dead cells. We used the “gold standard” of CFU to identify these conditions. The conditions containing all live cells were the 6-hour timepoints of 25°C, 30°C and 35°C for the temperature treatments and the 0-, 0.5-, 3-, and 6-hour timepoints of the 0% ethanol treatment. These are typically permissive conditions for yeast cell growth. The conditions containing mostly dead cells (as judged by CFU) were the 6-hour timepoints of 55°C and 65°C for the temperature treatments and the 6-hour timepoints of the 20% and 80% ethanol treatments. Both the 20% and 80% ethanol conditions were used as dead labels as we wanted to include extreme (80%) and milder conditions to induce death (20%).

The final time point of 0/5% ethanol was selected as LIVE labels and the final time point of 20%/80% ethanol was selected as DEAD. A similar labeling was provided with heat killed samples, namely the final time point of the first two and final time point of the last two gradients were selected as LIVE and DEAD, respectively. These were selected to provide the model with some very healthy, and most likely healthy cells, as well as very dead and dying/dead cells. The labels are called weak labels because the underlying hypothesis of the training regimen is that the majority of events will be labeled correctly, but it does not require all the events to be labeled correctly. In other words, we select control conditions that will have the expected majority of cells to be alive and dead. The reason we select the two highest gradients of treatment is to provide the model context with potentially different properties of dead - ones that either change or keep the cell’s morphology intact. A model is then trained with six channels as input (FSC-A, FSC-H, FSC-W, SSC-A, SSC-H, SSC-W) and is applied to all other conditions to provide all events with a preliminary set of labels. This model can be a neural network or any other machine learning model that can provide a set of preliminary labels to data from all other conditions. We chose a random forest classifier (RFC) for this first stage as it was significantly faster for training and inference. This model can be compared to stain based approaches of identifying live and dead cells.

#### Stage 2: Training Neural Network with Custom CFU-Based Objective

The second stage of training uses a fully-connected neural network model to update the labels based on a CFU measurement per condition (Figure 2D). Specifically, the objective function of the model can be divided into two parts:

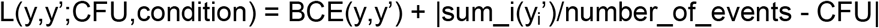

The first term seeks to reproduce the labels that were learned by the first stage model while the second term updates the weights based on context from the CFU from each condition. Given the objective selects a step per batch, the batch size for this model needs to be larger than what one would usually use (we used 2048 versus common batch sizes of 32 or 64). This ensures that each batch has a sufficient number of events per condition. We do not require each condition to be in the batch, but do want about 30 events from a condition in the batch. We found that setting the batch size to 2048 would ensure that. The model can now be used on any other data to annotate live and dead events.

### Evaluation of AutoGater with Held out Experiments

Given two stages of training and the abundance of data generated from a large corpus of experiments, we evaluated the model in two ways:

1. **Event based comparison with stained data:** Event based comparison of model to manually using a Sytox stain. In this test, the model was trained with data that was stained, but the color channel that measured the stain was not provided as a feature to the model for training. In this way, we could identify the events that the model predicts as live and dead and correlate those events with the stain color channel.
2. **Population based comparison with unstained data:** Another goal of the modeling framework was to harmonize the different methods in which cells are considered alive and dead. In this case, we trained a model with unstained data and compared percent live and dead to the same experimental conditions with stained and CFU data. In this way, we expect to see AutoGater’s predictions fall in between the stained and CFU percentages as the loss function jointly optimizes both terms.

As a reminder, the modeling framework presented selects a subset of conditions to label as *live* and *dead* (see Stage 1). A model is trained with those conditions and then evaluated on all held out conditions. The CFU measurements at each condition then updates the labels to harmonize the definition of live/viable cells across both stain based and CFU measurements. Conditions that measured CFU were never held out and are only used during training of the model. We should make clear that while the experiments were conducted over time for both heat and ethanol treated cells, time was never a feature that was input to the model. Thus, any temporal trends discovered by the model are an indirect artifact of the FSC and SSC features, as well as the CFU measurement at the point in time.

## Results

### Autogater Successfully Identifies Live/Dead Regions

For the event based comparison of cells, our goal was to identify if the model would be able to identify events that would be gated out by an expert as debris or dead cells. As a reminder, in this instance, the model was trained with data from cells that were stained with Sytox, but the color channel that measured Sytox was not used as a feature for the model. Figure 3A shows how an expert might select live and dead events choosing a threshold to divide unstained vs. Sytox stained cells. Autogaters predictions for live (Figure 3B) and dead events (Figure 3C) are also shown. We should note that the boundary line is not produced in the predictions, it is simply reproduced from the manually selected set of events for comparison. Over 90% of Autogater’s predictions of live events are on the left of the manually selected set of events. Interestingly, ∼60% of Autogater’s predictions of dead events are also to the left of the line. Recall that Autogater looks to balance both event based labels as well as CFUs for which we already saw a significant discrepancy in Figure 2C.

**Figure 3:**
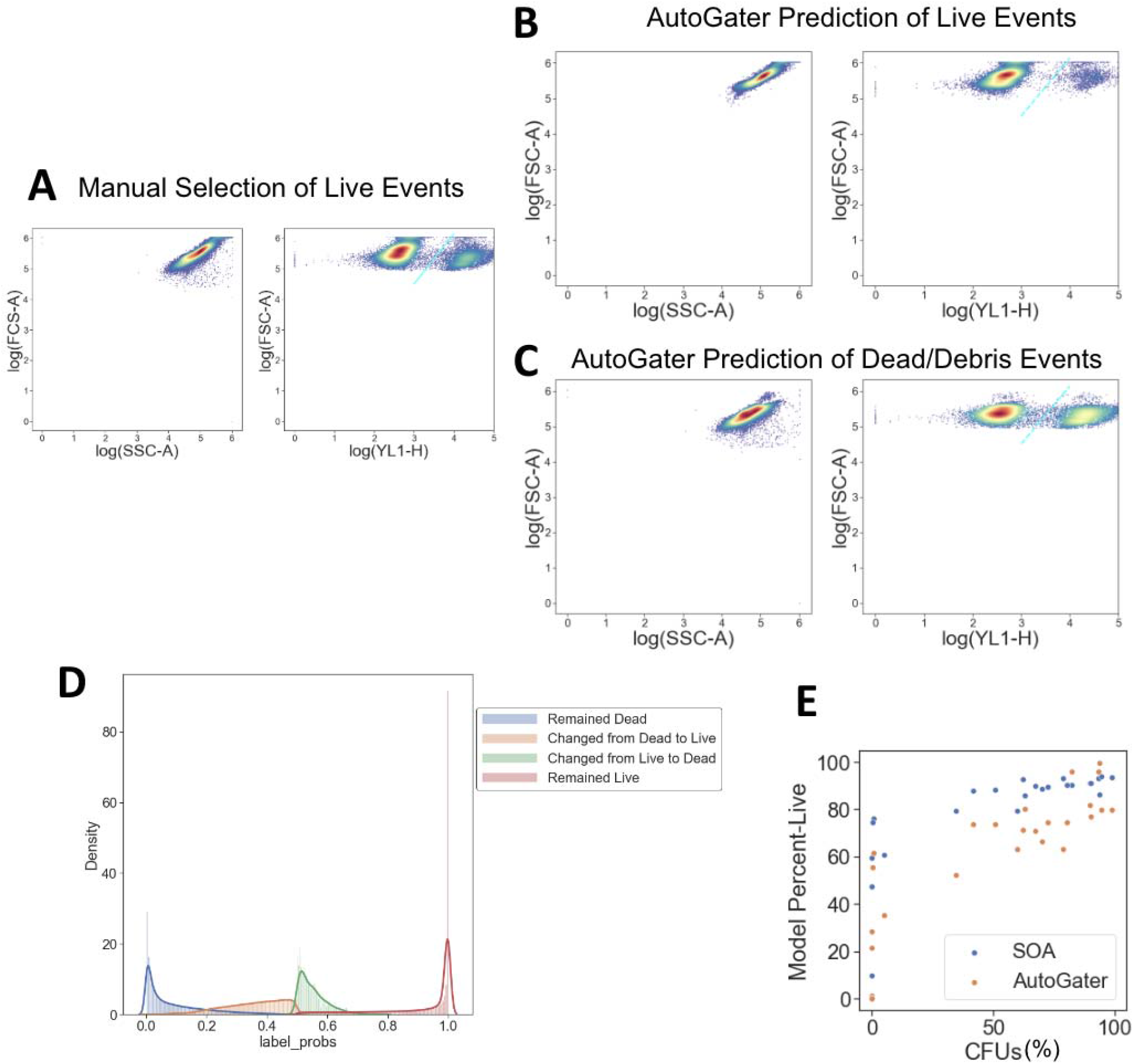
Characterization of model with Sytox data across all ethanol conditions. (A) Manual gating using Sytox channel of ethanol killed yeast cells to be used as a comparison for Autogater. (B) Testing the model on a sample it was not trained on shows Autogater is able to reconstruct the population of healthy, viable cells versus events that should be gated. The majority of the predicted live cells fall to the left of the gating line. Note: While showing the color channel, this channel was not provided as input to the model for its prediction. (C) Testing the model on a sample it was not trained on shows Autogater is able to reconstruct the population of dead cells versus events that should be gated. The population of dead cells to the right of the line is much larger than those of the live cells. The majority of cells however still fall to the left of the line, which means that Autogater is more conservative in what it calls live from the stain based method. Note: While showing the color channel, this channel was not provided as input to the model for its prediction. (D) Events that the model selected to update from live to dead and vice versa when incorporating the CFU measurements were ones that a model that did not use CFU information was least confident about (those with p∼0.5). (E) Using CFU as the ground-truth (x-axis), we compared the stain-based method (SOA) and AutoGater predictions to the CFUs and see that without using stains, AutoGater is more restrictive than stain-based methods to classify a cell as a live-cell. AutoGater’ predictions align better with CFUs at the lower and medium values of CFUs, but its restrictive nature gates out more events than needed at higher-CFUs.

Given the number of events classified as dead to the left of the line in Figure 2C, we wanted to see which events did the model think are dead but the stain would indicate they are alive. Recall, the second stage of the training framework refines the live/dead annotations by including CFU data while Stage 1 is trained on labels from the samples and does not include CFU. Therefore, it is not surprising that there are events to the left of the line that the model would consider dead. Figure 3D shows that most of Autogater’s changes (the orange and green curves) are events in which the first stage of Autogater, namely the ones that did not use information from CFUs, was unsure (probability of ∼0.5). This probability is output by the model and is a measure of the model’s live (probability of 1) and dead (probability of 0) prediction. We conclude from this analysis that the model uses CFUs to update the labels of events that it is most unsure of their live/dead state while still preserving the labels of its confident predictions.

To further measure the correlation between the manual selection (SOA), AutoGater, and CFUs, Figure 3E shows model predictions vs. CFU measurements. It is immediately clear that both Sytox and Autogater tend to underestimate the percentage of dead cells in the population as compared to CFU. This result could indicate that unhealthy or dying cells in the population that are not able to form colonies over time, are not identifiable as dead by Sytox or Autogater. That said, AutoGater better accounts for CFUs in the mid to large range, which indicates it assesses LIVE better than dead. The R^2^ between CFUs and Sytox stained vs AutoGater from Stage 2 is 0.05 versus 0.55, respectively. This finding demonstrates a significant impact on CFU information in AutoGater’s ability to identify dead events and harmonize event based and population based measurements of death. While not shown in the plot, we also measured the correlation between CFU and Autogater Stage 1 and Stage 2. Stage 1 only uses the weak labels while Stage 2 accounts for the CFU data. In this case, we observed an R^2^ of 0.4 and 0.55, a 37.5% increase in performance. The impact of the CFU data for training is noticeably evident and enables AutoGater to harmonize the two measurements by only using forward and side scatter channels.

### A focus on Ethanol Based Killing of Cells

Since different modes of cell death may produce different results, we explored multiple environmental stresses that induce cell death. We first examined ethanol stress at multiple concentrations over time. For all methods of screening live/dead cells in ethanol treated populations, a trend of increasing death occurs as ethanol concentrations are increased as well as treatment time increases (Figure 4 Panels A-G). Additionally, as the ethanol concentrations increase, the rate at which cells die increases.

**Figure 4:**
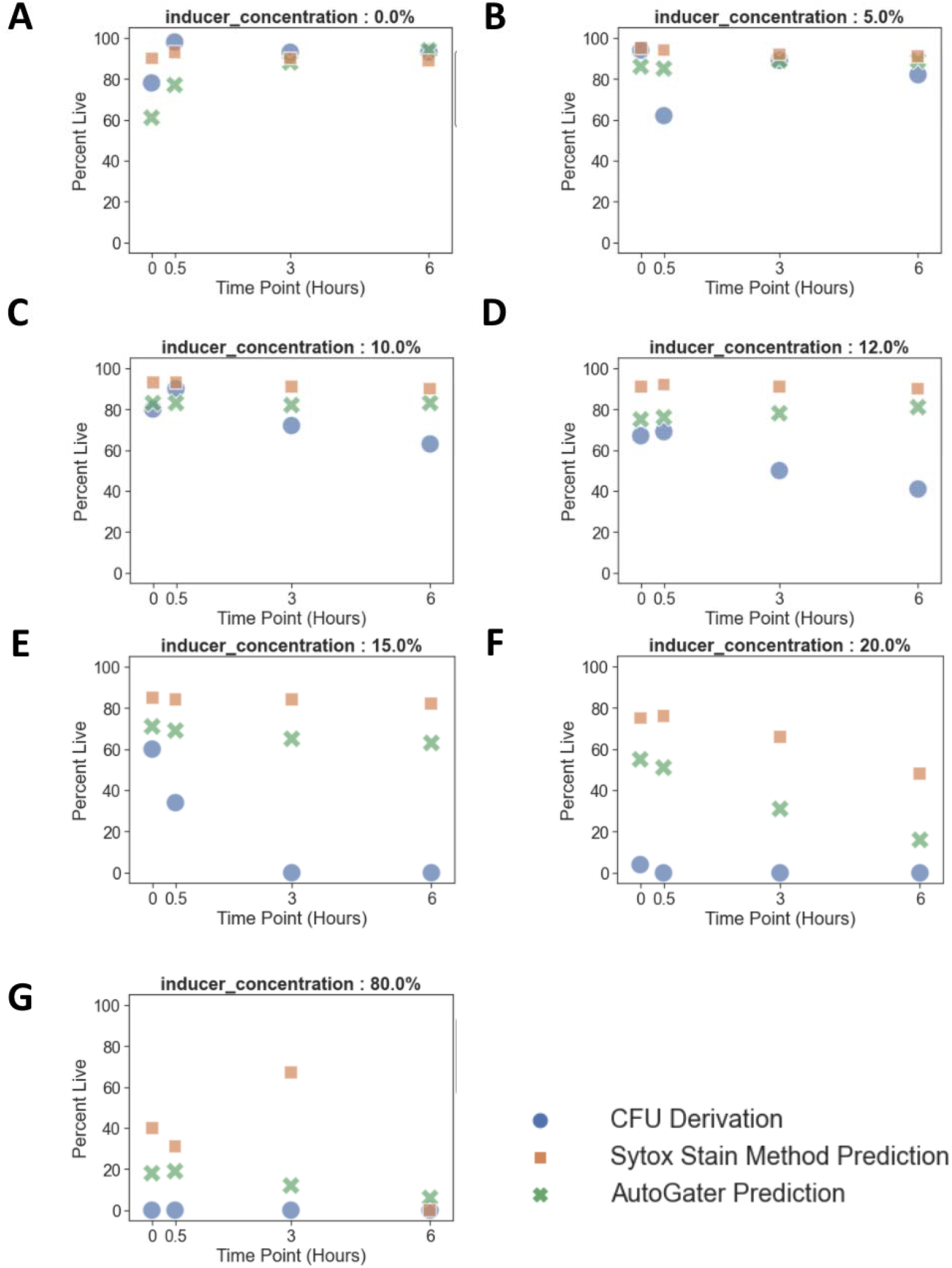
Comparison of various methods (colors and style) to measure dead over time (x/y axes) and ethanol concentrations (panels A-G). CFU derivations are population based measures of death, Sytox stained are cell based proxy measurements of death, and AutoGater are predictions from the neural network that use a weakly supervised approach to refine condition labels of death with those of CFUs. In this case, the live label was the final time point at 0.0% ethanol, while the death was the final time point at (20% and 80%) ethanol. AutoGater is able to learn a method to tag dead cells that combines the cell based measurements and population based measurements. This is best observed in panel (F).

However, the actual rate of death between methods does differ as the percent of cells determined to be alive by each method mostly differ for each sample. As ethanol concentrations increase, the CFU method is typically the first to respond, followed by AutoGater and then SYTOX. This phenomenon is most notable in ethanol concentrations 15% and 20%. The fact that AutoGater can capture the trends is interesting as AutoGater does not take in time as a feature and therefore has no concept of time. Additionally, AutoGater’s predictions on what cells are alive mostly fall in between SYTOX’s and CFU’s estimates of live cells. SYTOX identifies cells as dead at the time of the staining if their cell membranes are breached and pumps are no longer active, allowing the stain to access DNA, CFUs identifies cells as dead based on their ability to divide at some later time, and thus might identify cells that are destined for death that might not have suffered a breach in their cell membrane at the time of analysis. Thus, SYTOX may overestimate the percentage of live cells at the time of analysis with respect to CFUs. We speculate that Autogater is detecting features of cells that are dead and/or dying at the time of the analysis.

### A Focus on Heat Based Killing of Cells

All three methods for screening live/dead cells in temperature treated populations displayed similar trends in their estimates of live cells as their estimates of lives cells in ethanol treated cells. The trend being an increase in cell death as temperatures increased and as time increased. This trend is perhaps even more apparent in the temperature treated experiments as they are in the ethanol experiments (Figure 5).

**Figure 5:**
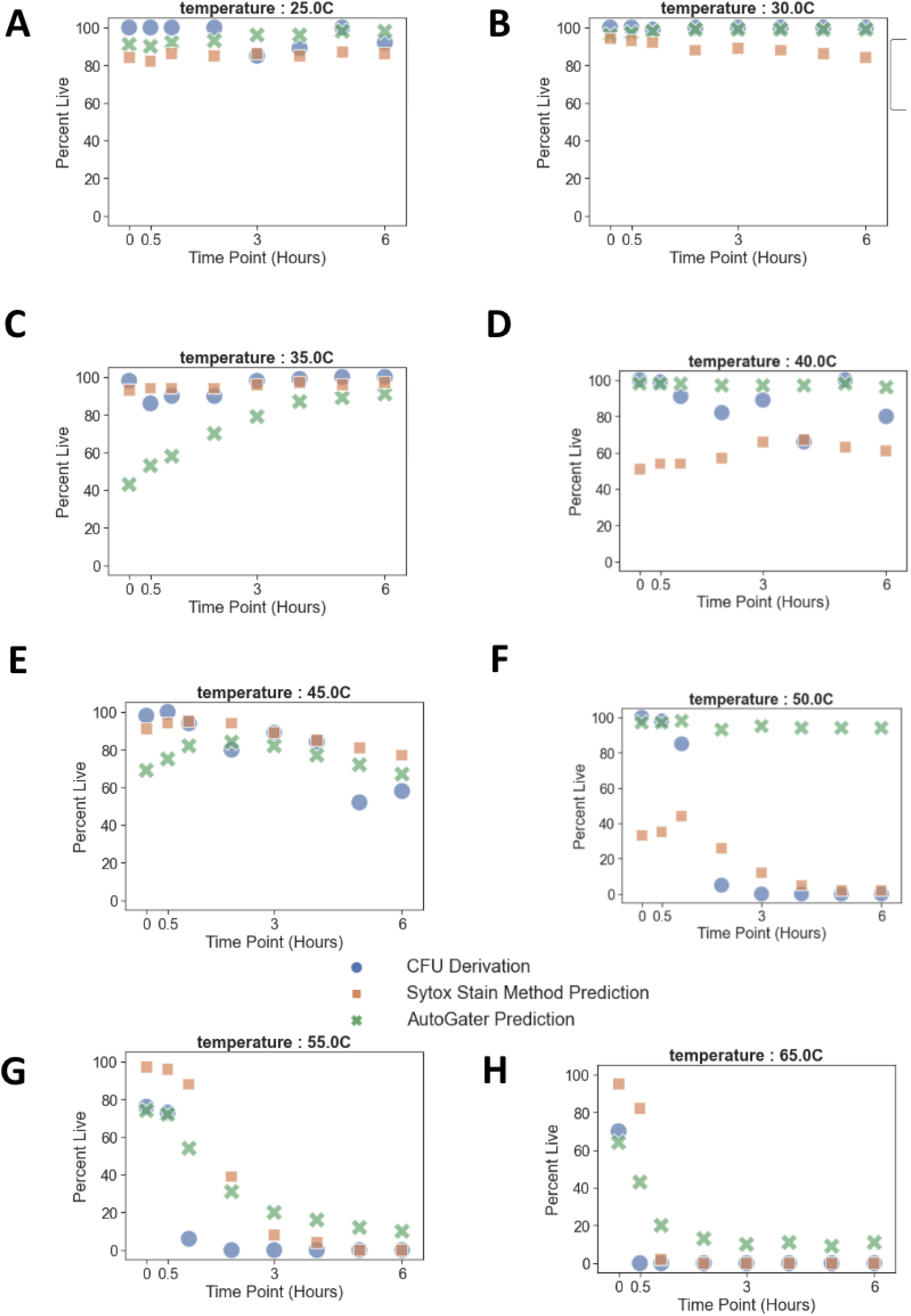
Comparison of various methods to measure dead over time and temperature. CFU derivations are population based measures of death, Sytox stained are cell based proxy measurements of death, and AutoGater are predictions from the neural network that use a weakly supervised approach to refine condition labels of death with those of CFUs. In this case, the live label was the final time point at 25° 30°, 35°C, while the death label was the final time point at 55° and 65°C. AutoGater is able to learn a method to tag dead cells that combines the cell based measurements and population based measurements. This is best observed in panel (F).

Additionally, AutoGater’s percent live predictions mostly fall in between STYOX’s and CFU’s percent live estimates as they did for the ethanol experiments. This suggests again that AutoGater is considering both the single cell and cell population perspectives of cell death. The samples treated at 40°C and 50°C seem to be an exception to this as CFU and SYTOX show an increase in cell death over time, however, AutoGater predicts an almost constant percentage of cells remaining alive over the course of both experiments. We did an analysis of the forward scatter and side scatter channels and noticed that for these two temperatures, the trend of a lower forward scatter at higher temperatures does not hold (Supplementary Figure 1). Given AutoGater only has context of forward and side scatter channels, if the phenotypic property is not captured in those channels then AutoGater will fail at its predictions. AutoGater learns and predicts live and dead cells using the FSC and SSC channels. Upon inspection of the data from these channels across all temperature treated samples, it was found that the samples from 40°C and 50°C had significantly large populations of events at the saturation point on the FSC channel. This observation led to filtering out all events at this saturation point and retraining AutoGater for subsequent predictions. AutoGater predictions for these two temperatures were not improved suggesting some other phenomena could be impacting the results.

## Discussion

Flow cytometry has revolutionized how we measure cellular phenotypes in disciplines extending from immunology to synthetic biology. A key capability in making these cell autonomous measurements is to eliminate dead cells that may confound the analysis. Eliminating dead cells is generally part of the preprocessing steps in the workflow where debris and doublets are also eliminated.

Current technologies for identifying dead cells take advantage of different features associated with dead or dying cells. Nucleic acid stains are commonly used and these only stain cells that have lost membrane integrity and/or the ability to pump these DNA stains out of the cell before they can stain DNA ^15,18,19^. Stains can also be used to identify cells that lack robust mitochondrial function. Finally, the microbiological approach for determining the percentage of dead cells in a population is to measure colony forming units (CFUs). CFU analyses identify the percentage of cells in the population that can undergo cell division in order to produce a colony.

Here we show that Autogater, using a neural network model, can utilize information across multiple channels to distinguish between live and dead cell populations in budding yeast cells. While the precise definition of dead cells utilized by Autogater is unknown, the model was trained on information only from Forward Scatter and Side Scatter channels. Using populations of dead and dying cells that were killed by either heat or ethanol treatment, we demonstrated that Autogater could identify dead cell populations that were similar, but not identical to cells stained with Sytox. Using training data from both SYTOX stained cells and CFU analyses, we found that Autogater estimates for dead cells tend to fall between SYTOX and CFU estimates (Fig. 4 and Fig. 5). Perhaps this finding is not surprising given that the model was trained on data from both SYTOX and CFU analyses.

With similar accuracy (Fig. 3) Autogater has a couple of significant advantages over nucleic acid stains or CFU analyses. The lack of a dependence on a stain saves money, sample preparation time and complexity as well as allowing the fluorescence channel to be used for other measurements. As well, staining approaches require the user to choose gates that separate live and dead cell populations. Finding an accurate gate is not always straightforward as the populations aren’t always well separated. Thus, user input on gating can hinder reproducibility, a substantial problem in flow cytometric analyses ^20,21^. Autogater provides an unbiased and automated method that should reduce costs and increase the speed and reproducibility of analyses. Finally, CFU analysis utilizes different features to identify dead cells. We found that CFU analyses tend to give higher estimates for the number of dead cells than SYTOX, suggesting that CFUs may more accurately identify cells that are in the process of dying at the time of analysis. That said, CFU analysis is a population measurement and cannot identify which cells in the flow cytometric analyses are dead or dying. When trained on both SYTOX and CFU analyses, Autogater appears to account for features of dead cells identified by both approaches while allowing real-time determination of which cells are dead or alive.

Currently, Autogater has been tuned to distinguish between live and dead cell populations in budding yeast only. We examined whether Autogater trained on yeast cells could work on bacterial cells, but had limited success. We tested the first stage of training for AutoGater across multiple single-cell organisms (*Escherichia coli* MG1655, *Bacillus subtilis* Marburg 168, and *S. cerevisiae*) and found the performance to have 50% accuracy, or random chance, across organisms (Supplementary Fig. 2). We do expect that the model could be trained on data collected from a variety of cells to learn features of dead cells specific for each cell type.

AutoGater’s two-stage training framework, namely the weakly supervised neural network approach, is also not limited to identifying healthy viable phenotypes. We envision this training framework to be extended to any application that needs single-cell based predictions for higher level phenotypes. The weak labels provide the model the ability to learn the high-dimensional patterns across multiple channels that make up the phenotype, a technique often reserved for experts to gate based on low-dimensional patterns, typically just on fluorescence and scatter channels. The joint objective function can also be extended to incorporate additional population based measures for the phenotype. Thus, we expect that AutoGater’s approach will be generalizable for assessing complex and phenotypes that are not directly observable in a measurement made on a stain for a particular molecule.

## Author Contributions

M.E, R. M, and S.H designed the experiments. R.M executed the experiments and collected the data. M.E and H.E designed and conducted all analyses and models. D.B provided initial guidance on the use of machine learning for the task. All authors contributed to manuscript development.

## Data Availability

All analyses and data used for this manuscript can be found here:

Ethanol:

https://github.com/SD2E/pysd2cat/blob/development/src/pysd2cat/analysis/live_dead_pipeline/AutoGater%20-%20Duke%20v2%20on%20Duke%20v2%20-%20Stain.ipynb

Heat:

https://github.com/SD2E/pysd2cat/blob/development/src/pysd2cat/analysis/live_dead_pipeline/AutoGater%20-%20Heat%20-%20No%20Stain.ipynb

