## Supplementary figures and images for "AutoGater: A Weakly Supervised Neural Network Model to Gate Cells in Flow Cytometric Analyses"

### Supplementary Figures

#### Slide 1
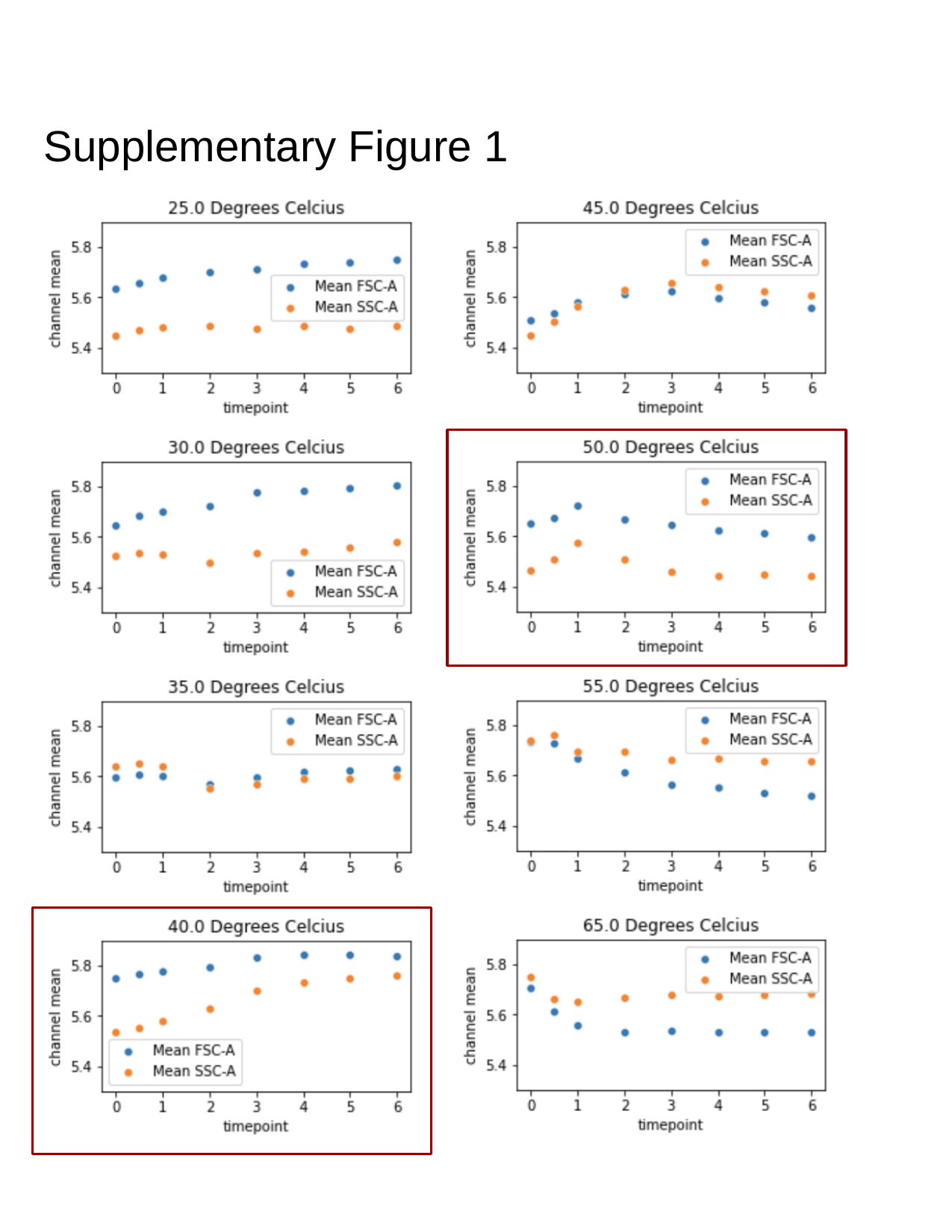

##### Supplementary Figure 1

#### Slide 2
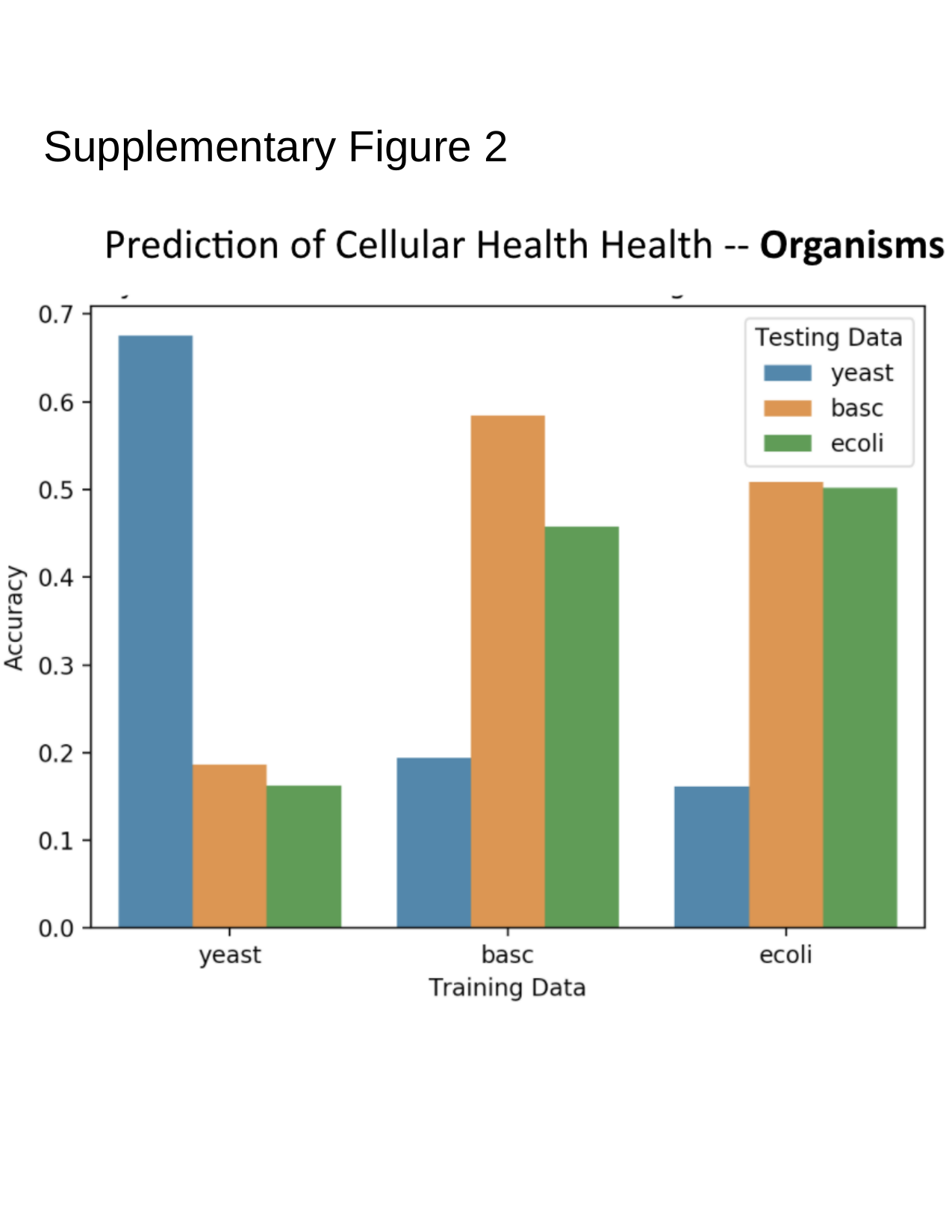

##### Supplementary Figure 2
